# Devising reliable and accurate epigenetic predictors: choosing the optimal computational solution

**DOI:** 10.1101/2023.10.13.562187

**Authors:** Charlotte D. Vavourakis, Chiara M. Herzog, Martin Widschwendter

## Abstract

Illumina DNA methylation arrays are frequently used for the discovery of methylation signatures associated with aging and disease. One of the major hurdles to overcome when training trait prediction models is the high dimensionality of the data, with the number of features (CpGs) greatly exceeding the typical number of samples assessed. In addition, most large-scale DNA methylation-based studies do not include replicate measurements for a given sample, making it impossible to estimate the degree of measurement uncertainty or the reliability of the prediction models. A recent study proposed that training penalized regression models on derived principal components (PCs) rather than on the original features (CpGs) results in more reliable age predictions, as estimated from technical replication. Moreover, the same method could be applied for predicting other phenotypes more reliably. Here, we aimed at validating the proposed PC method. We found that although dimension reduction with PCA consistently led to small improvements in the reliability of age prediction models, it severely compromised their accuracy. PC-based models needed far larger training set sizes to be similarly accurate as CpG-based models, whereas reliability did not depend on the sample size of the training set data for either approach. Finally, the PC version of a novel multiclass predictor for breast, ovarian and endometrial cancer we trained using weighted ensembles of deep-learning models also had a markedly lower predictive accuracy compared to a CpG version, suggesting limited applicability of the proposed PC method for predicting phenotypes beyond age.

## Introduction

The methylome is a flexible layer of the genome that plays a crucial role in the regulation of gene expression and chromatin structure. The strong correlation of aberrant DNA methylation patterns with various disease phenotypes can be used for training diagnostic tools with machine learning. For example, we previously developed the WID-BC [1], WID-OC [2], WID-EC [3] signatures that predict breast, ovarian and endometrial cancer from cervical smear samples analyzed using the Illumina Infinium-Methylation EPIC BeadChip v1.0, which covers approx. 850,000 CpG sites across the human genome. Because the methylome also changes with age in a predictable manner, another popular use for Illumina methylation array data is training age predictors, also called epigenetic clocks. Initially clocks were trained to predict only chronological age, i.e., the time from birth to a specific date [4–6]. The observation that deviations in actual versus predicted age correlate with certain phenotypic traits ranging from gender to mortality risk, sparked new DNA methylation-based investigations into biological aging, which is a measure of the age-related health status of an individual. Improved biological clock versions are now trained directly incorporating biomarkers for general health and physiological decline associated with age [7].

Two major obstacles present when training DNA methylation-based trait-prediction models. First, the microarray data are highly dimensional, with typically 100-or 1000-fold more CpGs than samples assessed. Secondly, in most large-scale DNA methylation-based studies, each sample is processed and measured only a single time. Because each measurement carries a certain degree of uncertainty which cannot be estimated without replication, continuous effort goes to understanding and reducing various sources of variability. Various data-driven pre-processing pipelines have been developed that remove noise, e.g. [8–10], and recently, Illumina released the MethylationEPIC v2.0 BeadChip, claiming improved precision by removing about 10% probes on the EPIC v1.0 that were deemed unreliable, at the same time adding probes to a total of 900K CpG sites to the array to better cover genomic regions relevant for clinical research [11].

Recently, Higgins-Chen *et al* [12] proposed that a simple dimension reduction done by training penalized regression models on extracted principal components rather than on the original features directly, results in more reliable age clocks because the PCA feature space incorporates information from all of the original features. Furthermore, the authors propose their procedure could be applied for training more reliable epigenetic biomarkers predicting other phenotypes than age. Here, we aimed at validating the proposed PC method by comparing both accuracy and reliability of CpG- and PC-based versions of the same clocks. Furthermore, we tested a use case for a PC-based cancer prediction model, by training a CpG- and PC-based version of a novel epigenetic, multi-class predictor for the detection of three different female cancers.

## Methods

### PC and PC proxy versions of the Hannum clock

Table 1 gives a summary of all datasets analyzed in this study. To investigate the performance of signatures retrained with the proposed PC approach [12], we first checked whether we could reproduce results for the Hannum clock using the same subset of 78,464 CpGs from the original train (“Hannum” [4]) and test (“BloodRep_450K”, GSE55763 [13]) datasets. Hannum age was calculated using the R package methyl-clock (version 1.2.1). Following the procedure in [12] and R code available from [14], PCs were used as features to train a regression model to predict chronological (PC approach) or the original Hannum age (PC proxy approach) with elastic net penalty and 10-fold cross validation to select the optimal tuning parameter. When using PCA as a dimension reduction step, the number of input features reduces to the number of samples in the training set (n) - 1. To calculate predicted age in the test set, methylation values for the test samples were projected on the trained PC space as described in [12]. Using the exact same procedure, we trained a new Hannum PC2 version on all 473,034 CpGs common between the train and all non-replicated samples available from GSE55763 [15], the latter “BloodFull_450K” used as a second independent validation set to compare the accuracy of the different clock versions. Missing values from GSE55763 were imputed using the R package impute [16] (version 1.68.0 27).

**Table 1.** Datasets analyzed in this study. Dataset ID = custom label used to denote each (composite) dataset; Type = type of Illumina DNA methylation array; n = final sample numbers carried forward for the analysis, excluding replication; Reps = number of technical replicates for each sample; Tissue = tissue from which DNA was analyzed; Repo = public repository from which the data was downloaded; Access ID = accession numbers used to access the data from the public repository.

| Dataset ID | Type | n | Reps | Tissue | Repo | Access ID |
| --- | --- | --- | --- | --- | --- | --- |
| repeatability | EPIC v1.0 | 4 | 4x | cervix | EGA | EGAS00001007184 |
| 3CDisc | EPIC v1.0 | 1657 | 1x | cervix | EGA | EGAS00001005055,<br>EGAS00001005045,<br>EGAS00001005033 |
| 3CExtVal | EPIC v1.0 | 449 | 1x | cervix | EGA | EGAS00001005055,<br>EGAS00001005045,<br>EGAS00001005033 |
| Hannum | 450K | 656 | 1x | blood | GEO | GSE40279 |
| BloodRep_450K | 450K | 36 | 2x | blood | GEO | GSE55763 |
| BloodFull_450K | 450K | 2639 | 1x | blood | GEO | GSE55763 |

### Quality control and pre-processing of MethylationEPIC data sets

We analyzed three data sets generated with the Illumina MethylationEPIC BeadChip v1.0, starting from the raw data publicly available in IDAT format under restricted access (Table 1). All data sets were pre-processed as described previously (1). Detection of failed probes (detection p-value *>* 0.01) was performed with the R package minfi [8] (version 1.40.0). Samples with median methylated or unmethylated intensity *<* 9.5 or with *>* 10% failed probes were removed. Background intensity correction and dye bias correction was performed with the preprocessNoob function in minfi. Probe bias correction was performed using the beta mixture quantile normalisation (BMIQ) algorithm in the R package Champ [9] (version 2.24.0). Probes that mapped to the Y chromosome, SNPs flagged by [1, 8, 17], non-CpG probes, and probes that failed in *>* 10% of the samples were removed. Missing values were imputed using the R package impute [16] (version 1.68.0 27).

### Effect of training size on prediction model performance

To test the effects dimension of reduction by PCA on age prediction model performance as a function of training size, we used the same procedure to train penalized regression models on chronological age using sample subsets of increasing size from the “450K_BloodFull” (n min = 34, n max = 2639; 473,864 CpGs) and “3CDisc” data (n min = 32, n max = 1657; 776,725 CpGs). Per experiment, Elastic net regression models were trained for each random sample subset once directly on all CpGs and once on derived principal components (n-1) as input features. Each experiment was performed three times, recording in each pass for the trained model’s accuracy estimates (slope of the regression predicted versus actual age) calculated from the “3CExtVal” set and reliability (intraclass correlation coefficient; ICC) estimates calculated from the “repeatability” set [18]. Results from the three experiments were combined and power law curves were fitted to the scores obtained for each model using nonlinear least squares with the nls() nls.control() functions in the R package stat (version 4.2.0):

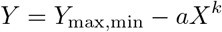

with *Y* = the performance metric and *X* = the number of samples in the trained (sub)set, *Y*_max,min_ = the maximum or minimum metric the model converges to and a,k the shape parameters of the power law curve.

### PC and non-PC versions of a multi-class cancer classifier trained with AutoGluon

The “3CDisc” set with 1647 subjects in the age range 19-95 years at time of consent (869 healthy women [5], 329 breast cancer [1], 242 ovarian [2] and 217 endometrial cancer cases [3]) was used to train different versions of a multi-class classifier predicting breast, ovarian or endometrial cancer with the AutoML framework AutoGluon-Tabular [19] (version 0.6.1-cpu-jupyter-ubuntu20.04-py3.8, Docker Hub digest ace9e4217b43). With this framework, data processing, deep learning and model ensembling with k-fold cross validation (bagging) is performed in an automated way within a user-specified time constraint. Ensembles are made using a multi-layer stacking strategy, in which the predictions of n base learners are concatenated with the original data features in each k out-of-fold bag for training higher-layer stackers that ultimately are weighted and aggregated into an ensemble. The “3CExtVal” set with 449 subjects in the age range 24-86 years at time of consent (225 healthy women [5], 113 breast cancer [1], 47 ovarian [2] and 64 endometrial cancer cases [3]) was used for testing accuracy of the classifiers.

Before training, we removed a set of probes flagged internally to have a consistently low normalized, mean fluorescence intensity and disregarded data from those probes that did not make it to Illumina MethylationEPIC BeadChip v2.0, meaning that the same analysis can be performed with the latest available version of the EPIC array once EPIC v1.0 has been discontinued. Predictions of cell composition were done on the pre-processed beta-values using published algorithms. The immune cell fraction in each sample was estimated using the EpiDISH R package (version 2.12.0), using the epithelial, fibroblast and immune cell reference dataset “centEpiFibiC.m” [20].

For the non-PC version “multi-3C-autogluon” of the classifier, an initial feature selection step was introduced because the first data preparation step in a trial with 632,385 features (number CpGs left after initial pre-processing) did not complete within a reasonable time (48 hours). An initial selected set of features was used for a first training round with 2:3 of the training data (original train:test split used by [1–3]). The remaining 1:3 of the “3CDisc set” was used to test the importance of each feature for the predicted label by the best weighted ensemble model and only features with an importance score *≥* 0 were used for a second, final training round on the full “3CDisc set”. For the first training round, the features with the top 5000 absolute delta-beta values for the immune and/or epithelial component for each individual cancer in the 2:3 training subset were selected as described in [1]. Delta-beta values were calculated by linearly regressing the beta values for cases and controls separately for each feature (CpG) against estimated immune cell composition and age, and calculating differences of the intercept points at immune cell proportion = 0 or 1. Initial feature selection resulted in 17,863 input CpGs from which only 28 were removed before the second training round.

For the PC version “multi-3C-autogluon PC” of the classifier, PCA was performed with the sklearn.decomposition.PCA class of the Python scikit-learn package (version 1.1.3), calling pca.fit transform() on the training and pca.transform() on the test data sets. To deal with missing features in the test data sets, we calculated the beta-value with maximum density across all samples in the training set and used this as a fixed value in the test set for the respective missing feature. For both the non-PC and the PC version, chronological age of each subject and estimated immune cell composition of a sample (EpiDish) were inputted as additional features, which were retained for the second training round for the non-PC version. For all signatures, the TabularPredictor was trained using the preset = ‘best quality’ for fit(), with a time constrained of 3,600 s for the initial training round of the non-PC version, and of 10,800 s in all final training rounds.

### Accuracy and reliability estimates

To estimate accuracy (i.e., the closeness of the predicted values with the true values) of the age clocks, root mean squared errors (RMSE) were calculated for linear regressions of predicted versus actual chronological age in the test data sets. To compare the accuracy of the Hannum clock with the PC2 version clock, the “BloodFull_450K” dataset was subsetted to 479 blood samples from individuals in the same age range 24-75 years as those in the training set “Hannum”. To estimate reliability (i.e., the closeness of repeated predicted values) of the trained age prediction models, ICCs were calculated with the R package irr (version 0.84.1; CRAN) for the test datasets comprised of technical replicates using single rater, absolute-agreements and two way randomeffects [12]. For the multi-class cancer classifiers, accuracy was estimated by the AUC from multiclass ROC curves constructed with prediction probabilities testing each test class versus the rest. Confusion matrices and AUC scores were calculated using the sklearn.metrics.confusion matrix() and sklearn.metrics.roc auc score() functions called from the Python scikit-learn package (version 1.1.3).

## Results and discussion

First, we verified that we could reproduce the PCA approach and age predictions from [12]. PC and PC proxy versions of the Hannum clock [4] were retrained to predict chronological age and Hannum age, respectively, on the original subset of 78,464 CpGs from the HumanMethylation450 dataset derived from 656 peripheral blood samples in the dataset “Hannum” (see Table 1 for a summary on the datasets analyzed in this study). Age predictions from the resulting clocks on a blood-derived test set comprising 36 duplicate measurements “450K_BloodRep” were then compared to resulting age predictions reported by [12]. We obtained the same predicted values for the Hannum clock in the test set “BloodRep_450K” as [12] (Fig. 1, panel a). A small offset was found between the Hannum PC proxy predicted ages (slope and intercept of the regression = 1 and -0,22, Fig. 1, panel b), but this difference is expected and negligible considering that during training the folds for cross validation are chosen randomly.

**Fig. 1.**
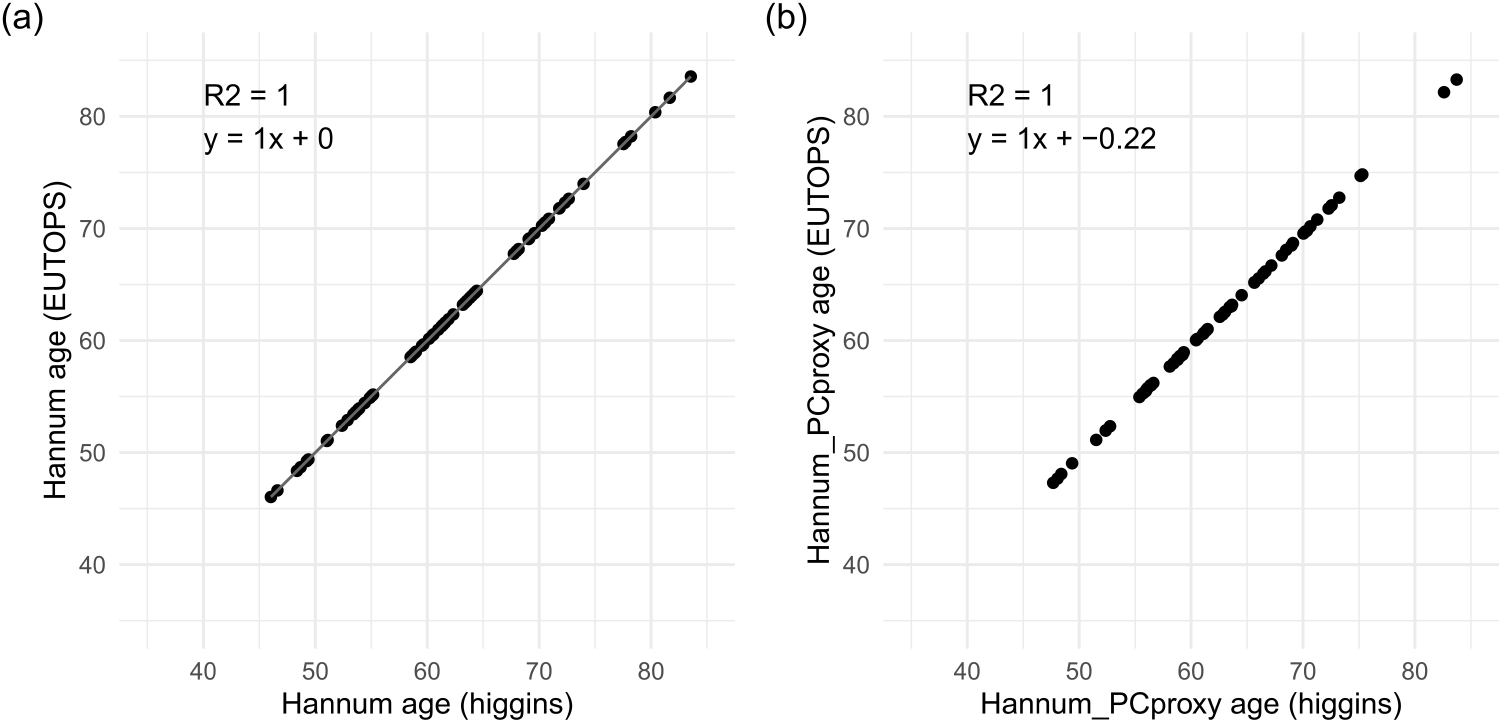
Reproducing clock calculations from [12] Regression of age predictions for the same clock versions calculated ourselves (Y-axis) versus by [12] (X-axis) for (a) the original Hannum clock and (b) the PC proxy version of the Hannum clock

As reported by [12], reliability as estimated by the Intraclass Correlation Coefficient (ICC) improved for the predictions of the PC versus the original Hannum clock version calculated for the test set “BloodRep_450K”. However, age predictions from the PC clock were significantly less accurate, with a 19% lower root-mean-squared-error (RMSE; Fig. 2, panel (a)) and a significantly lower regression slope for predicted versus actual age (slope = 0.661, 95% CI: 0.556-0.767 for the Hannum PC clock and 0.855, 95% CI: 0.771-0.940 for the Hannum clock; Fig. 2, panels (b-c)). This loss in accuracy was not reported by [12]. Training a second PC clock on a larger subset of 473,034 CpGs shared between the “Hannum” training set and the test set “450K_BloodFull” (n = 2639 blood samples) aggravated the loss of accuracy, with a regression slope of only 0.59 (95% CI 0.57 - 0.60) for the Hannum PC2 clock (Fig. 3). This suggests that the loss of accuracy of training clocks in the PCA feature space might be a general issue.

**Fig. 2.**
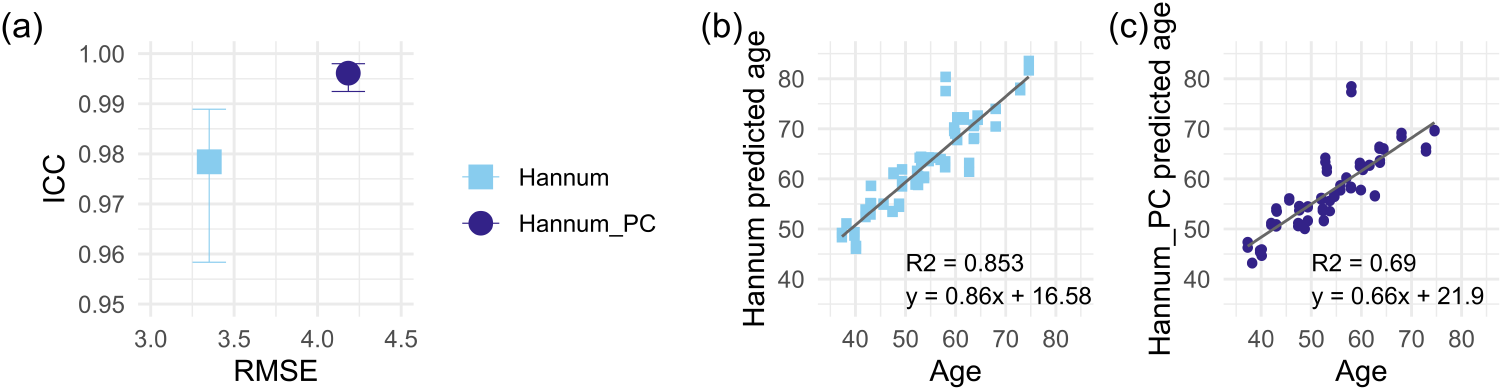
Performance of original and PC versions of the Hannum clock [12] in the test set “Blood-Rep 450K”. (a) Precision (ICC; error bars represent 95% confidence intervals) and accuracy (RMSE) of the linear regression between predicted age versus chronological age, with corresponding regressions for (b) the original Hannum clock and (c) the PC version of the Hannum clock. The PC and PC proxy versions of the Hannum clock were trained on the 450K “Hannum” dataset (*n* = 656 samples, 78,464 features used) to predict age and Hannum age, respectively, reproducing the method proposed in [12]. Age predictions were done for 2 *×* 36 technical replicates from blood samples in the “Blood-Rep 450K” dataset.

**Fig. 3.**
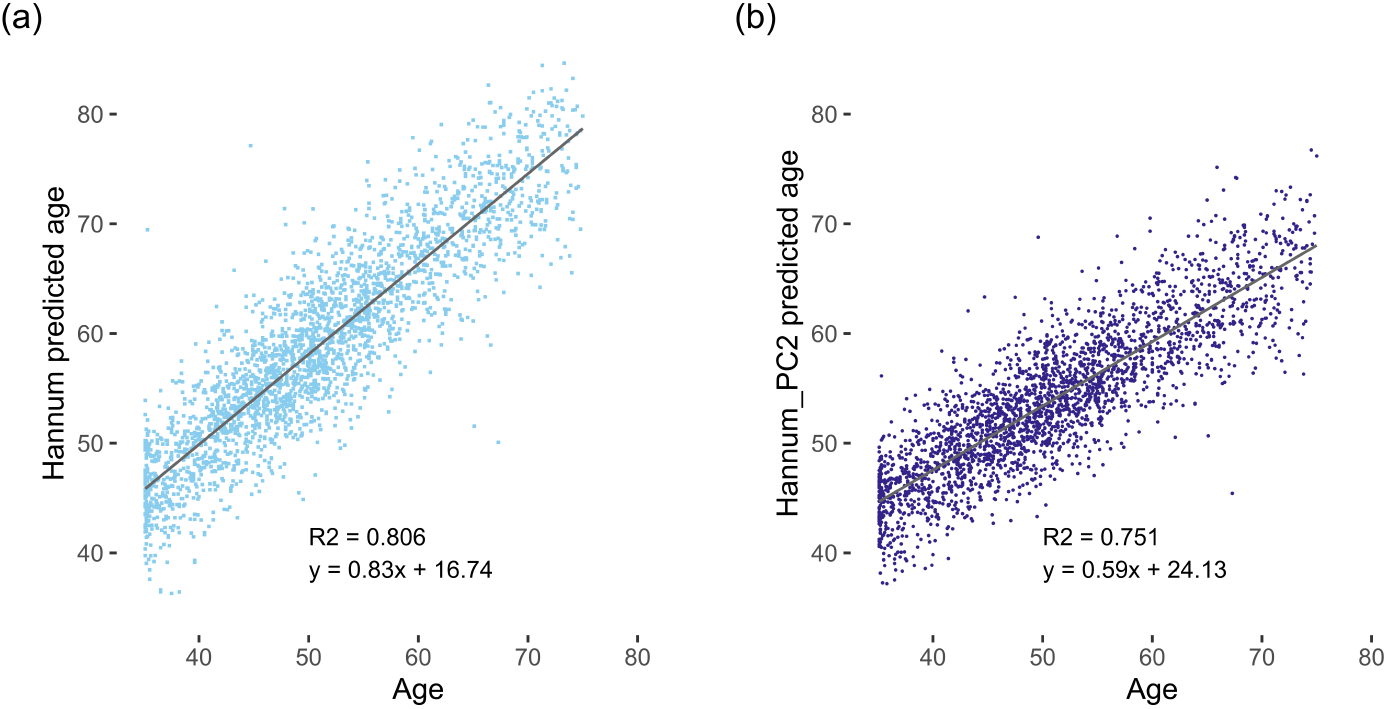
Accuracy of the original and PC2 versions of the Hannum clock in the test set “Blood-Full 450K”. The PC2 Hannum clock version was trained on the 450K “Hannum” dataset (n=656 samples age range 24-75 years, 473,034 features used) to predict age using the method proposed in [12]. Age predictions were done for non-replicated blood samples in the “BloodFull_450K” dataset from individuals in the age range 24-75 years (n=2639 blood samples).

Next, we investigated how the observed tradeoff between accuracy and reliability of age prediction models underlying clocks directly depends on the additional PCA step and the size of the training set. Increasing, randomly fixed sample subsets of the “450K_BloodFull” (n = 2639 blood samples) and the EPIC v1.0 data set “3CDisc” (n = 1647 cervical smear samples) were used to train Elastic net regression models in the original or the PCA-derived feature space, recording for each training round accuracy and reliability of age predictions in external validation sets of corresponding sample and data types (Fig. 4). The same experiment was repeated three times and power-law curves were fitted to the model performance estimates. Over the complete size range of training subsets, the Elastic net regression models had a much lower predictive accuracy when trained on PCs instead of CpGs for both the blood (Fig. 4a) and cervical smear DNA methylation data (Fig. 4c). Whereas the clocks trained on the original feature space neared the modeled maximum accuracy for the full training sets (Ymax = 0.90*±*0.05 for blood and Ymax = 0.94*±*0.04 for cervical smears in the modeled fits), hypothetically the PC clocks would reach a comparable accuracy of 0.9 only for approximately 11,707 blood or 17,000 cervical smear samples (calculated from the modeled equations). In contrast, differences in reliability were minimal between PC and non-PC clocks, mostly showing an excellent ICC well above 0.9 [21] (Fig. 4b/d). Only for the cervical smear data, few individual clocks showed random drops in ICC, something the PC-trained clocks did not suffer from (Fig. 4d). Thus, Fig. 4 clearly demonstrates that training penalized regression models on the PCA-derived feature space can come with a considerable loss in accuracy and that the accuracy of PC clocks strongly depends on the sample size of the training set.

**Fig. 4.**
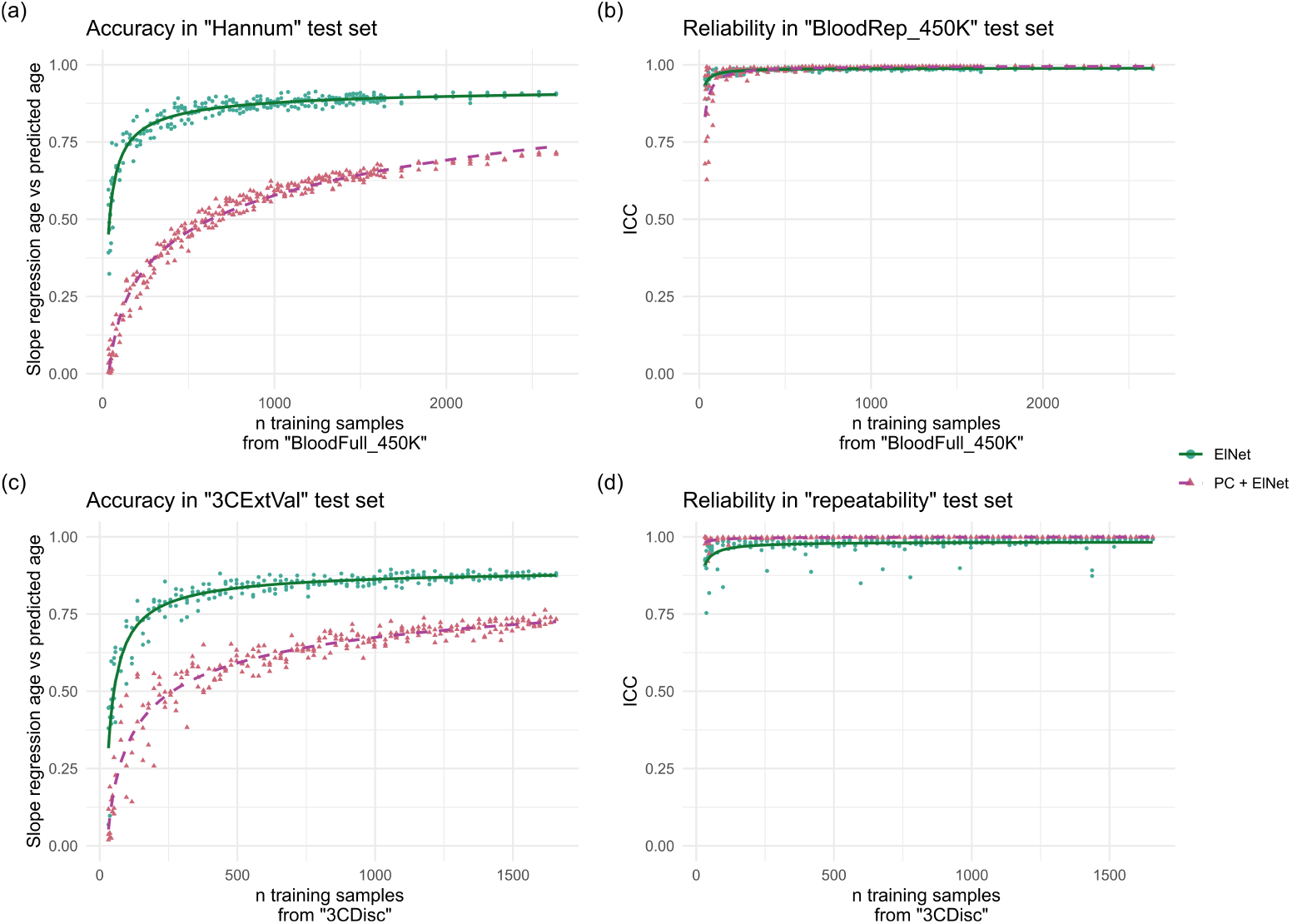
Performance of Elastic net regression models trained as a function of training size directly on the same set of CpGs as input features (Elnet, green shades and point shapes) or on n-1 derived PCs (PC, purple shades and triangle shapes). Performance parameters of each clock from three independent experiments (points) and modeled fits (lines; Y-axis) are plotted as a function of n samples in the training set (X-axis), with n max = 2639 for the blood-derived **(a-b)** and n max = 1647 for the cervical-smear derived DNA methylation dataset **(c-d). (a)** Predictive accuracy of the age clocks in blood (n = 656 samples). Elnet fit: *Y* = 0.94(*±*0.04) *−* 4.5(*±*0.6)*X*^*−*0.63*±*0.04^; PC fit: *Y* = 6.94(*±*0.02) *−* 8(*±*4)*X*^*−*0.03*±*0.02^; *±SE*. **(b)** Test-retest reliability of the corresponding clocks (n = 36 samples, 2 x measured each). Elnet fit: *Y* = 0.99(*±*0.08) *−* 1.5(*±*0.5)*X*^*−*0.92*±*0.08^; PC fit: *Y* = 1.0(*±*0.2) *−* 7(*±*4)*X*^*−*1.1*±*0.2^; *±SE*. **(c)** Predictive accuracy of the age clocks in cervical smears (n = 449 samples). Elnet fit: *Y* = 0.90(*±*0.05) *−* 9(*±*2)*X*^*−*0.79*±*0.05^; PC fit: *Y* = 1.07(*±*0.03) *−* 2.6(*±*0.2)*X*^*−*0.28*±*0.03^; *±SE*. **(d)** Test-retest reliability of the corresponding clocks (n = 4 samples, 4 x measured each). Elnet fit: *Y* = 1.0(*±*0.3) *−* 2(*±*2)*X*^*−*1.0*±*0.3^; PC fit: *Y* = 1.0(*±*0.2) *−* 0.2(*±*0.2)*X*^*−*0.7*±*0.2^; *±SE*.

The variability in age-related health outcomes can be interpreted as an interindividual difference in biological age relative to chronological age [7]. The deviation between chronological age and predicted age using epigenetic biomarkers has previously been used as an estimate of biological age and has been shown to predict multiple agerelated health outcomes, such as mortality. Interestingly, Higgins-Chen *et al* [12] report a stronger association of some versions of their PC clocks compared to the corresponding non-PC clocks. For instance, PCHorvath1 and PCPhenoAge exhibit a stronger association with mortality than Horvath1 and PhenoAge, respectively. Another recent DNA methylation-based study also found a stronger correlation between predicted and actual telomer length when transforming the data with PCA before training [22]. However, in agreement with our models (Fig. 4), Zhang *et al* [6] previously demonstrated that highly accurate chronological age predictions from blood samples are feasible when the sample set size used for training is large enough, with error rates compared to chronological age approximating zero. As the association of a clock with chronological age increases, the association with health outcomes shrinks, and a trained epigenetic biomarker then only informs on chronological age. Many recent epigenetic biomarkers are now training to predict biological outcomes rather than chronological age alone to overcome this limitation [7]. Although Higgins-Chen *et al*. [12] demonstrate increased performance for predicting age-related outcomes beyond the initial outcome variable of chronological age with many of their PC clocks, our data provide additional context and reveal a loss of accuracy when training penalized regression models on PCA-transformed DNA methylation data, particularly for chronological age - the key outcome variable these clocks are designed to predict.

To further demonstrate how dimension reduction by PCA may affect prediction model accuracy in the context of a directly-linked clinical outcome, we compared the accuracy of different versions of a cancer prediction model trained on an EPIC v1.0 dataset derived from 1647 cervical smear samples from healthy women [5], age-matched with breast (BC) [1], ovarian (OC) [2] and endometrial cancer (EC) [3] cases (Fig. 5). Simple penalized regression models were replaced by ensembles of rich deeplearning base models trained with the AutoGluon framework [19] to allow for multiclass predictions. The accuracy of the classifier trained on the original feature space was comparable to those of individual cancer prediction models trained with simple penalized regression models (AUC WID-BC = 0.81, 95% CI: 0.76–0.86 [1], AUC WID-EC = 0.92, 95% CI: 0.88-0.97 [3], AUC WID-OC = 0.76, 95% CI: 0.68–0.84 [2]; Fig. 5a). Meanwhile, the PC version of the classifier suffered from a lower accuracy, with areas under the receiver operator curve (AUC) dropping from 0.82 to 0.66 for predicting BC, from 0.93 to 0.89 for predicting EC, and from 0.70 to 0.55 for predicting OC, respectively (Fig. 5b).

**Fig. 5.**
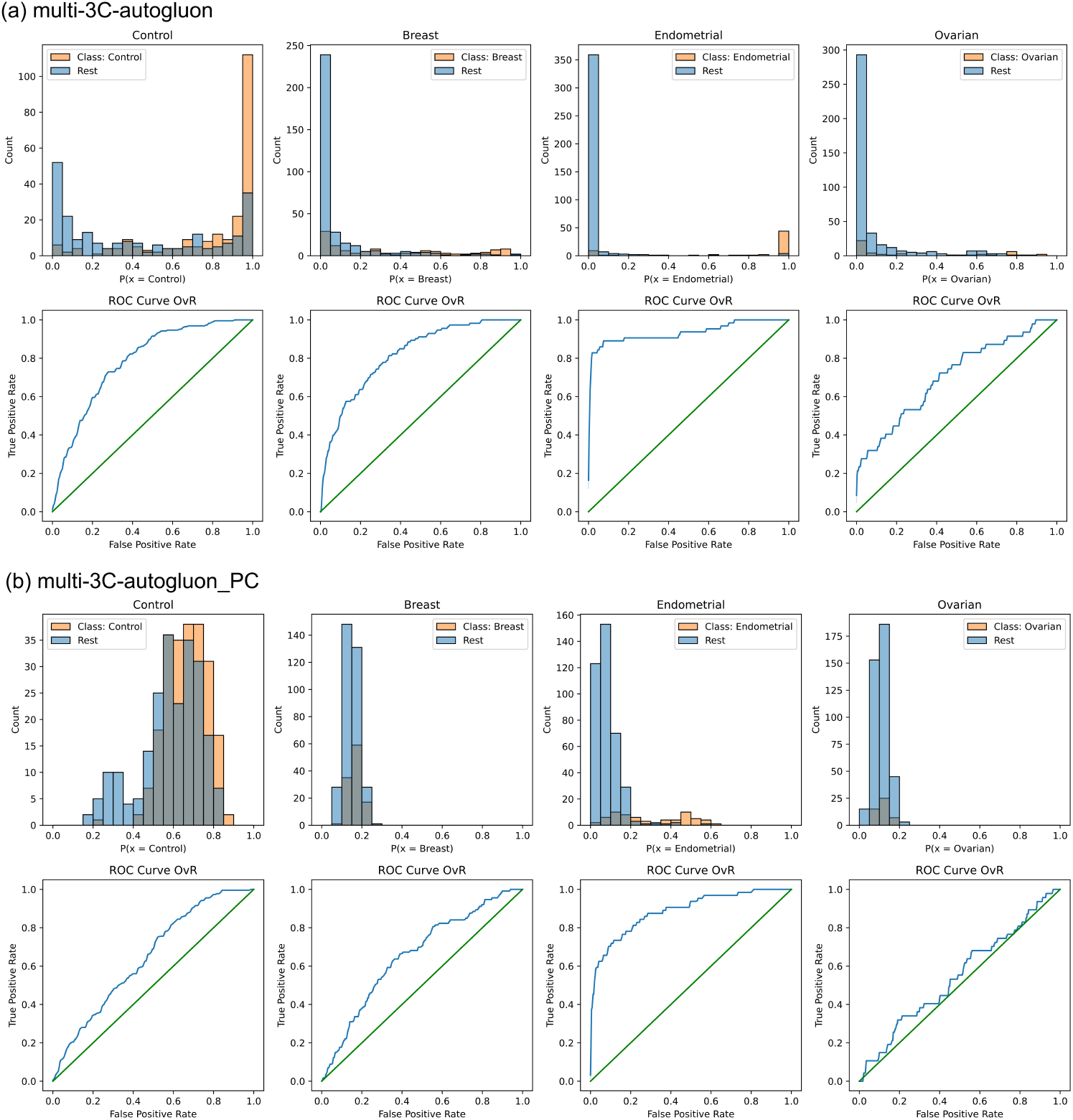
Effect of feature engineering on the performance multi-class cancer prediction models. (a) “multi-3C-autogluon” is a version of the classifier trained on 17,837 selected features from the original feature space (beta-methylation values of CpGs). (b) “multi-3C-autogluon PC” is a version of the classifier trained on 1646 principal components derived from the beta-methylation values, as well as (chronological) age of a sample’s subject and predicted immune cell composition of a sample. Multiclass classifiers were trained on the “3CDisc” data with the automated machine learning framework AutoGluon and accuracy was evaluated on the “3CExtVal” data comparing each class versus the rest of the classes. Top rows in each panel show the distribution of the predicted probabilities for controls (cancer free women), breast, ovarian or endometrial cancer cases. Bottom rows in each panel give the corresponding receiver operator (ROC) curves.

Some important technical considerations should be noted when comparing the performance of PC-based and CpG-based age prediction models as a function of the size of the training set, as we have done here. Firstly, when combining methylation data from an increasing number of samples, the chance that a given target CpG successfully yields a methylation readout in all samples decreases, and the resulting elimination of features that automatically occurs when combining methylation data from larger sample sets was not accounted for in the above model projections. It may positively affect performance when unreliable features are eliminated, but negatively affect performance when important features are lost. Secondly, for the CpG-based models the number of input features were fixed, hence for any n samples the same number of features (CpGs) are evaluated during training. In contrast, by design each PC-based model is trained on n-1 PCs, thus there is a direct dependence of the number of input features (PCs) on the sample size of the training set. Thirdly, the modeled Ymax values exceed 1 for the PC-approach. The modeled curves describe the recorded accuracies well in the tested sample set size range, but the extrapolated path is nonetheless hard to predict. Hypothetically, increasing the number of samples and hence the number of PCs could lead to overfitting, which can be overcome again either by increasing training size even more or by increasing model complexity [23]. Increasing model complexity by using state-of-the-art deep-learning approaches instead of simple penalized regression models such as Elastic Net further opens the potential for training multimodal trait prediction models that may evaluate methylation data, various omics data, biomarker data, and medical images at once.

## Conclusion

In summary, the optimal machine learning approach depends on the available data and intended purpose of the prediction model. Various approaches should be considered before settling on a final machine learning strategy. Although we could confirm that dimension reduction by PCA leads to more reliable age predictions, we conclude that dimension reduction by PCA is far less accurate compared with the same models trained in the original feature space, while the improvement in reliability is marginal. Therefore, the method proposed by [12] to improve reliability might not be suitable when fewer than 10,000 samples are used for training new epigenetic biomarkers. Although we do not expect that reliability of models trained in the original feature space is much poorer compared to PC-models, preferably a few technical replicates should still be included in any study design to check reliability in addition to accuracy. Finally, in the case of cancer prediction, we demonstrate here how an alternative means of reducing dimensionality by feature selection can lead to more accurate classification models compared with models trained on PCs, and further prove that the use of deep learning algorithms through AutoML platforms can be a simple way of leveraging the utility of DNA methylation array data.

## List of abbreviations

AUC: Area Under the ROC curve
BC: breast cancer
CI: confidence interval
CpG: Cytosine-phosphate-Guanine
EC: endometrial cancer
ICC: intraclass correlation coefficient
OC: ovarian cancer
PC: principal component
PCA: principal component analysis
ROC: Receiver Operating Characteristic

## Declarations

## Ethics approval and consent to participate

Not applicable.

## Consent for publication

Not applicable.

## Availability of data and materials

The 450K microarray datasets analyzed during the current study are available in the NCBI’s Gene Expression Omnibus and are accessible through GEO Series Accession Numbers GSE40279 and GSE55763. The EPIC microarray datasets analyzed during the current study are available under restricted access through the European Genome-phenome Archive (EGA), which is hosted by the EBI and the CRG, under the accession numbers EGAS00001007184, EGAS00001005055, EGAS00001005045, EGAS00001005033. All of the analyses were performed with publicly available R packages (R version 4.2.0) and Python libraries (Python version 3.8.13, supplied within the Docker Hub digest ace9e4217b43 for the framework AutoGluon-Tabular) using custom scripts that are made publicly available on https://github.com/ChVav/EPICPCsignatures.

## Authors’ contributions

The work reported in the paper has been performed by the authors, unless clearly specified in the text. Methodology, Software, Formal Analysis, Validation, Visualization, Writing - original draft preparation: C.D.V.; Writing - Review & Editing: C.D.V., C.H., M.W.; Conceptualization, Data curation, Supervision and Project Administration: C.D.V., C.H.; Funding Acquisition: M.W.

## Acknowledgments

This project has received funding from the European Union’s Horizon2020 research and innovation programme under grant agreement No 874662 (HEAP); the European Research Council Proof of Concept grant No. 101113534 (BRCA-PREVENT). The Eve Appeal (https://eveappeal.org.uk/); and the LandTirol. Views and opinions expressed are however those of the author(s) only and do not necessarily reflect those of the European Union. Neither the European Union nor the granting authority can be held responsible for them.

